# rDNAmine: A New Tool for the Analysis of Long Repetitive Sequences

**DOI:** 10.64898/2026.02.24.707673

**Authors:** Agnieszka Czarnocka-Cieciura, Natalia Gumińska

## Abstract

In this study, we introduce a novel approach for analysing long, repetitive genomic sequences. Our methods significantly advance research on rDNA polymorphism. First, we describe a technique for isolating high-molecular-weight DNA from individual chromosomes, enabling selective enrichment of sequencing libraries for extensive genomic regions of interest. Second, we present rDNAmine, a bioinformatic toolkit for capturing and examining large repetitive arrays in Oxford Nanopore sequencing data. This approach facilitates the study of polymorphisms within long repeats, bypassing traditional alignment-based methods and providing a more efficient and scalable solution for investigating repetitive regions. We demonstrate the effectiveness of our approach through the analysis of rDNA arrays in two yeast species, *Saccharomyces cerevisiae* and *Candida albicans*. In *S. cerevisiae*, rDNA arrays show limited polymorphism, while in *C. albicans*, we observe substantial variation in rDNA module size, with two distinct repeat populations within the array. These findings reveal species-specific differences in the structural organisation of rDNA *loci*, highlighting the diverse nature of tandem repeat architecture. The rDNAmine toolkit is broadly applicable to various organisms and repetitive genomic contexts, offering a versatile platform for studying repetitive sequences.

**Take Away:**

- Yeast rDNA serves as a benchmark to validate tools for analysing long repetitive sequences.
- A chromosome-specific DNA extraction method has been introduced to enable targeted enrichment of repetitive *loci*.
- The rDNAmine pipeline was designed to analyse long tandem repeats from noisy long-read data without requiring global alignment.

## Introduction

The ribosome is a large ribonucleoprotein complex that translates genetic information encoded in nucleic acids into functional proteins. The catalytic sites of ribosomes reside within an evolutionarily conserved core composed exclusively of ribosomal RNA (rRNA). Ribosome polymorphism, which can manifest as variation in rRNA sequence or protein composition, represents an understudied layer of post-transcriptional gene regulation. In *Saccharomyces cerevisiae*, changes in ribosomal protein composition have been observed during the transition from fermentable to non-fermentable carbon sources (Sun *et al*., 2021). In plants, ribosome populations influence the expression of specific groups of transcripts, driving proteomic changes in differentiating cells. Nevertheless, relatively little is known about the role of rRNA-polymorphic ribosomes in shaping eukaryotic proteomes.

Ribosomal RNA can vary in nucleotide sequence, as well as in the presence and localisation of specific chemical modifications (Sharma i Entian, 2022). The most extensive studies on RNA-polymorphic ribosomes have been conducted in human cells. The absence of a single modification, 1-methyl-3-α-amino-α-carboxyl-propyl pseudouridine at position 1248 (U) of 18S rRNA, has been observed in up to 45.9% of patients with colorectal carcinoma and is prevalent in 22 cancer types (Babaian *et al*., 2020). Alterations in the chemical modification patterns of rRNA are a potential target for novel cancer therapies (Cui *et al*., 2024).

Intraspecies polymorphism in ribosomal RNA-coding genes (rDNA) has been documented in numerous eukaryotic species. In the genomes of mice and humans, both intragenomic and intraspecies rDNA variants have been identified. RNA sequencing of rRNA isolated from polysomes have confirmed the presence of polymorphisms in rRNA molecules within translating ribosomes (Fan *et al*., 2022). However, studying the functional aspects of RNA-polymorphic ribosomes remains challenging. Most studies rely on short-read sequencing data, which constrains the identification of polymorphisms coexisting within a single repeat. Furthermore, ribosomes are large complexes containing both rigid and highly mobile elements within their tertiary structures, limiting the study of the ribosomal core to crystallographic and cryo-electron microscopy (cryoEM) techniques. Chemical modifications of rRNA also restrict the applicability of RNA sequencing methods. Investigating the diversity of rRNA-coding genes is further complicated by the structure of their respective *loci*. In Eukarya, these genes occur in multiple copies, forming arrays of several dozen to several hundred repeats (Symonová, 2019). Moreover, many organisms possess more than one rDNA *locus* (Smirnov *et al*., 2021).

Currently, tools for assembling long repetitive sequences are lacking. Even with long-read sequencing technologies, assembling rDNA *loci* sequences remains unfeasible due to the high similarity between repeating modules. Many bioinformatic methods for polymorphism analysis are also limited by the length of rDNA repeats. In *S. cerevisiae*, a single module containing the complete set of rRNA-encoding genes and regulatory sequences spans approximately 9,100 base pairs (Arnau *et al*., 2022). This length increases with organismal complexity, exceeding 11,000 base pairs in higher eukaryotes. Consequently, the reference genomes of most eukaryotic organisms contain only a single consensus sequence of rDNA repeats, which also impedes research on polymorphic ribosomes (Ganley i Kobayashi, 2007).

In this study, we present novel tools for analysing repetitive sequences characterised by long tandem repeats. We have developed a method for isolating DNA from specific chromosomes, facilitating research on rDNA polymorphism within their respective *loci*. Additionally, we have devised an efficient approach for retrieving long repeat sequences from reads obtained through Direct DNA Sequencing by Oxford Nanopore Technologies (ONT) (Wang *et al*., 2021). ONT long-read sequencing has become a valuable tool for various biological applications, offering affordability, real-time analysis, portability, and scalability, with continuous improvements in accuracy and yield (Scarano *et al*., 2024). An expanding range of bioinformatics tools further enhances its utility (Smith *et al*., 2020; Krawczyk *et al*., 2024; Gumińska *et al*., 2025). Although the accuracy of ONT reads continues to improve, it remains lower than that of Illumina reads. Nonetheless, certain applications can achieve comparable accuracy while addressing some of Illumina’s limitations, particularly by increasing assembly contiguity in repetitive and low-complexity regions (Carbo *et al*., 2023; Linde *et al*., 2023; Szoboszlay *et al*., 2023; Zhang *et al*., 2023; de Bruin *et al*., 2025; Perlas *et al*., 2025).

The tabular data format generated by our software is straightforward to implement and process within the R or other programmatic environments. This approach facilitates the investigation of polymorphisms within individual repeats. Moreover, analysing rDNA copy sequences directly from reads provides valuable insights into the organisation of the rDNA array and variability in module size. In this study, rDNA serves as a well-characterised model system to benchmark our enrichment procedure and computational pipeline on inherently noisy Oxford Nanopore datasets. Our primary aim is to demonstrate that both the experimental workflow and the rDNAmine algorithm perform reliably. Here, we do not seek to address specific biological questions but rather to provide validated tools for downstream investigations.

## Materials and Methods

### Cell culture

A single yeast colony was cultured in 25 mL YPD medium overnight at 30°C (*S. cerevisiae* strain S288C) or 37°C (*C. albicans* strain BWP17) with shaking at 220 rpm. The seed culture was then inoculated into 300 mL shake flasks containing 50 mL fresh YPD medium with an initial OD 600 of 0.01 and cultivated at 30°C/37°C and 220 rpm until the OD reached 0.4.

### DNA isolation protocol

Yeast cultures were centrifuged at 3,000 rpm for 3 min at 4°C. The medium was removed, and the cells were resuspended in 25 ml of autoclaved water at 4°C. This step was repeated twice. Agarose blocks containing cells were then prepared according to the protocol (Herschleb, Ananiev i Schwartz, 2007).

The blocks were embedded in an agarose gel made with 0.8% High Resolution Plus agarose (BioShop) and 1% TAE buffer. PFGE gels were separated using a Pulsed Field Gel Electrophoresis Systems CHEF mapper (BioRad) in 1% TAE buffer at 18°C. The gel was then rinsed in ethidium bromide solution and photographed under blue laser light using a Typhoon™ laser-scanner (Cytiva). Based on the life-size image, sections containing the target chromosomes (for *S. cerevisiae* chromosome XII and for *C. albicans* chromosome R) were excised from the gel and placed in an electroelution chamber (BioRad). Electroelution was carried out for 10 h at 21°C.

The DNA-containing solution was transferred with a Pasteur pipette to an Amicon Ultra-2 Centrifugal Filter 50 kDa MWCO Millipore column (Merck). The sample was centrifuged at 3,000 rpm for 5 min at 4°C to remove most of the solution. Then 2 ml of ultra-pure water (Sigma) was added to the column and gently rocked for 1 min, followed by centrifugation at 3,000 rpm at 4°C for 5 min. This washing step was repeated twice. After washing, the sample was concentrated by centrifugation at 3,000 rpm at 4°C, with the duration adjusted based on the DNA sample’s density. The concentrated DNA was then transferred to Eppendorf tubes and stored at 4°C for up to one month.

DNA material prepared in this manner is highly sensitive to shaking and should not be frozen before library preparation. When transferring DNA samples, pipettes with trimmed tips or Pasteur pipettes are recommended to minimise nucleic acid fragmentation.

### Library preparation and sequencing

Isolated DNA samples were used to construct DNA-seq libraries starting with 400–500 ng of DNA. DNA were fragmented using DNase I (ThermoFisher Scientific) according to manufacturer’s protocol. RNA-seq libraries were prepared using the Ion Total RNA-Seq Kit v2 (ThermoFisher Scientific) following the manufacturer’s protocol beginning from the cDNA amplification step.

The libraries were sequenced on the Ion Torrent Proton™ NGS System according to the manufacturer’s instructions. Raw sequencing data were preprocessed using Torrent Suite™ Software (Life Technologies). Barcode removal and quality trimming were performed in Torrent Suite™ using default parameters. The processed reads were then exported as FASTQ files (sample accession numbers: ERS22921397 for *S. cerevisiae* and ERS22921399 for *C. albicans*).

PacBio library preparation and sequencing were outsourced to the Museum and Institute of Zoology Polish Academy of Sciences in Warsaw (sample accession numbers: ERS22921398 for *S. cerevisiae* and ERS22921400 for *C. albicans*).

### Mapping and quantification of sequencing depth

Sequencing reads obtained for *S. cerevisiae* were first mapped to the complete reference genome S288C_reference_genome_R9-1-1 (https://sgd-prod-upload.s3.amazonaws.com/S000216130/S288C_reference_genome_R9-1-1_19990210.tgz) from the Saccharomyces Genome Database (SGD). *C. albicans* sequencing reads were mapped to the reference genome ASM18296v3 (https://www.ncbi.nlm.nih.gov/datasets/genome/GCF_000182965.3/) from the National Center for Biotechnology Information (NCBI).

Mapping was performed using minimap2 with the following settings: -ax splice -uf -k 14 -t 20 (Li, 2018) for IonTorrent data and -ax map-pb -t 20 for SMRT PacBio data. Sequencing depth information was extracted from the BAM files using the samtools depth -a command (Danecek *et al*., 2021). SMRT and IonTorrent sequencing datasets were mapped and further processed independently. This mapping confirmed the enrichment of each sequencing dataset with chromosome sequences isolated during DNA sample preparation. Sequencing depth was then used to estimate the number of reads corresponding to each position on chromosome XII for *S. cerevisiae* and chromosome R for *C. albicans*.

### *De-novo* assembly of rDNA arrays

Long reads were aligned to themselves using minimap2 -x ava-pb -t 20. Assembly from the overlap graph was generated using miniasm –f (Li, 2016). Thehe assembly file was then polished with short reads using minimap2 -ax sr function.

### Estimation of rDNA copy number

To estimate the rDNA copy number, we used an IonTorrent dataset generated for *S. cerevisiae*. All reads were mapped to the rDNA reference module sequence using minimap2 with the following parameters: -ax splice -uf -k 14 -t 20 (Li, 2018). Sequencing depth information was extracted from the resulting BAM files using the samtools depth -a command (Danecek *et al*., 2021). All reads were then mapped to the entire reference genome (S288C_reference_genome_R9-1-1) from the SGD, and the overall sequencing depth was estimated. The average sequencing depth for chromosome XII of *S. cerevisiae* was calculated, excluding the region containing rDNA sequences (both repeats and pseudogenes). The average sequencing depth for reads containing rDNA and mapped to the rDNA reference module was also determined. The rDNA copy number was estimated by dividing the average sequencing depth calculated for rDNA by the average depth for chromosome XII. This approach was expected to provide an accurate estimate of the rDNA copy number in the genome.

### Nanopore direct DNA sequencing datasets used for the analysis of rDNA module polymorphism

ONT data used in this study were retrieved from the Sequence Read Archive (SRA) repository. Dataset SRR13441294 (https://www.ncbi.nlm.nih.gov/sra/?term=SRR13441294) for *C. albicans* (*Microbiology Resource Announcements*, 2021) was generated using the Direct DNA Sequencing method R9 with a MinION device (ONT). The run contained 412,984 reads with a total size of 4.9 G bases. This dataset was selected because more than half of the reads exceeded 11,000 bp in length. Dataset SRR17374240 (https://www.ncbi.nlm.nih.gov/sra/?term=SRR17374240) for *S. cerevisiae* (Zhang *et al*., 2022) was generated using the Direct DNA Sequencing method with a PromethION device (ONT). The run contained 701,791 reads with a total size of 9.6 G bases. This dataset was chosen because more than half of the reads exceeded 10,000 bp in length (Mak *et al*., 2023).

When selecting datasets for studies involving long repetitive sequences such as rDNA, we followed two key principles. First, the datasets needed to be enriched with reads exceeding twice the length of the rDNA repeat for the species in question (sought length intervals (in bp) for *C. albicans*: min = 11,000, max = 12,568; for *S. cerevisiae*: min = 9,000, max = 9,254). At the initial stage of our ONT data analysis, all reads shorter than 50,000 bp were filtered out to avoid issues related to pseudogenes, as discussed in detail in the second part of the Results section. Second, ensuring a minimum read length exceeding two full rDNA repeats guaranteed that at least one complete copy was present in the reads selected based on the presence of rRNA-coding sequences.

### Estimation of lengths of rDNA copies

The information about the rDNA module length was taken from FASTA files containing repeats selected using the rDNAmine tools with the following command:

awk ‘/^>/ {if (seqlen) print header, seqlen; header=substr($0, 2); seqlen=0} !/^>/ {seqlen += length($0)} END {if (seqlen) print header, seqlen}’ *.fasta *.fa | awk -v OFS=’,’ ‘{print $1, $2}’ >> module_length_summary.csv

### Determination and analysis of sequences of long repetitive modules

We have developed rDNAmine, a toolkit for analysing rDNA repeats based on data from ONT Direct DNA Sequencing. The input for the analysis is FASTQ files. The first stage of the pipeline is executed by the rDNAmine_miner.sh programme, which converts the FASTQ file into a set of FASTA files, each containing an individual read. Subsequently, rDNAmine_prospector.sh uses profile Hidden Markov Models (pHMMs) to identify reads containing repeat sequences. This step requires a FASTA file with the reference sequence of a single repeat as input. Using the HMMER algorithm (Finn, Clements i Eddy, 2011) (nhmmer --tblout $hmm_outtab), the read sequences are searched for similarity to the reference module sequences. The module coordinates obtained from the HMMER output files are then used to extract the rDNA modules from the reads.

The isolation of modules is performed by the rDNAmine_module_collector.sh programme, which filters out modules shorter than the specified cut-off (option -d sets the minimum expected module size, and option -l sets the maximum expected module size). This filtering is essential to prevent the analysis of incomplete rDNA copies. Next, the rDNAmine_collection_inventory.sh programme compares the module sequences with the reference module using the Multiple Alignment using Fast Fourier Transform (MAFFT) G-INS-i (Global Iterative Alignment using the Needleman-Wunsch algorithm with iterative refinement) (Katoh i Standley, 2013). Pairs of compared module-reference sequences are then converted into a tabular format by the rDNAmine_mine_format.sh program. In this format, the module sequences are analysed within the R environment.

### Analysis of module diversity

The mine.csv files, generated by the rDNAmine_mine_format.sh programme, were loaded into the R environment as tabular data. These files were then merged into a consolidated table, which served as the basis for further analyses.

To calculate the substitution-deletion polymorphism coefficient (SDC), table entries containing deletions and substitutions were used. The SDC provides a straightforward estimate of how frequently a given trait (not limited to nucleotides, but any observable trait) appears in the dataset. For each column in the table, representing a nucleotide position in the reference, we calculated the frequencies of each nucleotide and deletions (represented as ‘-’). These frequencies were then divided by the number of modules used in the analysis. The distribution of this coefficient was visualised using ggplot2 (Wickham, 2016). This coefficient describes the diversity of modules isolated from long reads obtained through Nanopore sequencing, rather than the direct polymorphism of rDNA repeats in the matrix located on chromosome XII of the yeast from which the DNA material was isolated. It is important to note that the value of the coefficient depends on two main parameters. The first is the accuracy of sequencing, which in Nanopore technology is gradually improving. The second is the type of algorithm used to perform alignments between the reference module and the modules isolated from the reads. Depending on the approach used, the alignments may contain more or fewer mismatches and may vary in length.

Next, a custom-prepared table, rDNAmine_S_cerevisiae.csv, was loaded into the R environment. This table contains information about the positions of genes and intergenic regions within the reference module, active sites of the ribosome, its tertiary structure, and phenotypes of mutations described in the literature. Using this information, modules containing more than one mutation causing a deleterious phenotype were filtered out from the combined rDNAmine_S_cerevisiae.csv and result tables.

In the reference-module tables generated by the rDNAmine_mine_format_generator.sh programme, insertions are marked within the reference position numbering by appending letters to the numerical position identifiers. This allowed us to filter the consolidated table, created by merging the mine.csv files, to isolate only those module positions containing insertions, along with information on the specific reference positions flanking insertions. We then estimated the maximum insertion lengths for each pair of adjacent positions in the reference sequence. The resulting data were visualised as the maximum insertion length for each reference position.

When aligning sequences that have been pre-screened using an HMM based on similarity, MAFFT can refine the alignment by incorporating structural and evolutionary constraints. The pre-screening step ensures that only homologous sequences are considered, improving the accuracy of the final alignment. This iterative refinement strategy further enhances alignment for phylogenetic and functional analyses.

### Comparison of *C. albicans* long and short rDNA module reference sequences

Module sequence analysis was performed using the Assemblytics tool (Nattestad i Schatz, 2016). Code is available on GitHub (https://github.com/MariaNattestad/Assemblytics).

### Data visualisation

Data provided by rDNAmine toolkit were further processed in R 4.4.1 (R Core Team) using base R and dplyr (Wickham *et al*., 2022) functions for sorting, filtering and pivoting, and the graphics were generated using ggplot2 library (Wickham, 2016).

### Hardware and software

Computations were performed on a Unix machine (22.04.4 LTS (Jammy Jellyfish), Intel Xeon Platinum E5-2683 v4 ×64 CPU 2.10 GHz, 64 cores, 512 GB RAM) with the following third-party software: EMBOSS 6.6.0 (Rice, Longden i Bleasby, 2000), HMMER 3.3.2 (Finn, Clements i Eddy, 2011), MAFFT 7.525 (Katoh i Standley, 2013), Miniasm-0.3 (r179) (Li, 2016), minimap2 2.28 (Li, 2018), samtools 1.20 (Danecek *et al*., 2021), seqtk 1.4 (Li *et al*., 2013), R 4.4.1 (Race for Your Life, R Core Team) with the following packages: dplyr 1.1.4 (Wickham *et al*., 2022), ggplot2 3.5.1 (Wickham, 2016).

rDNAmine was also tested on a desktop computer (Linux Mint 21.3 (Virginia), Intel Core i5-10210U ×64 CPU 1.60 GHz, 8 cores, 32 GB RAM) with the same software versions installed as stated above. The pipeline was designed for broad use on various configurations with minimal software dependencies. Processing time will be proportional to hardware capabilities.

### Data and code availability

All sequencing datasets were deposited in the European Nucleotide Archive (ENA) (https://www.ebi.ac.uk/ena/browser/home) under accession numbers: ERS22921397, ERS22921398, ERS22921399 and ERS22921400.

All scripts used for the analysis and the rDNAmine_S_cerevisiae.csv table can be found in the GitHub repository https://github.com/LRB-IIMCB/rDNA_mine.

## Results and Discussion

### Enrichment of DNA samples with material derived from specific chromosomes

To study rDNA clusters, we developed a procedure for selective DNA isolation (Figure 1A). The resulting biological material (Supplementary Figure 1A) was then sequenced using two independent methods: IonTorrent and SMRT long-read sequencing. In both organisms studied, the sequencing datasets were enriched with chromosome sequences containing rDNA arrays (Figure 1B-D). Due to the small differences in the sizes of chromosomes IV and XII in *S. cerevisiae* (Supplementary Figure 1B) and chromosomes I and R in *C. albicans* (Supplementary Figure 1C), accurately separating these chromosomes proved difficult, even with PFGE. However, in the species studied, this was not an issue as only one chromosome in each pair contained the rDNA repeat arrays.

**Figure 1.**
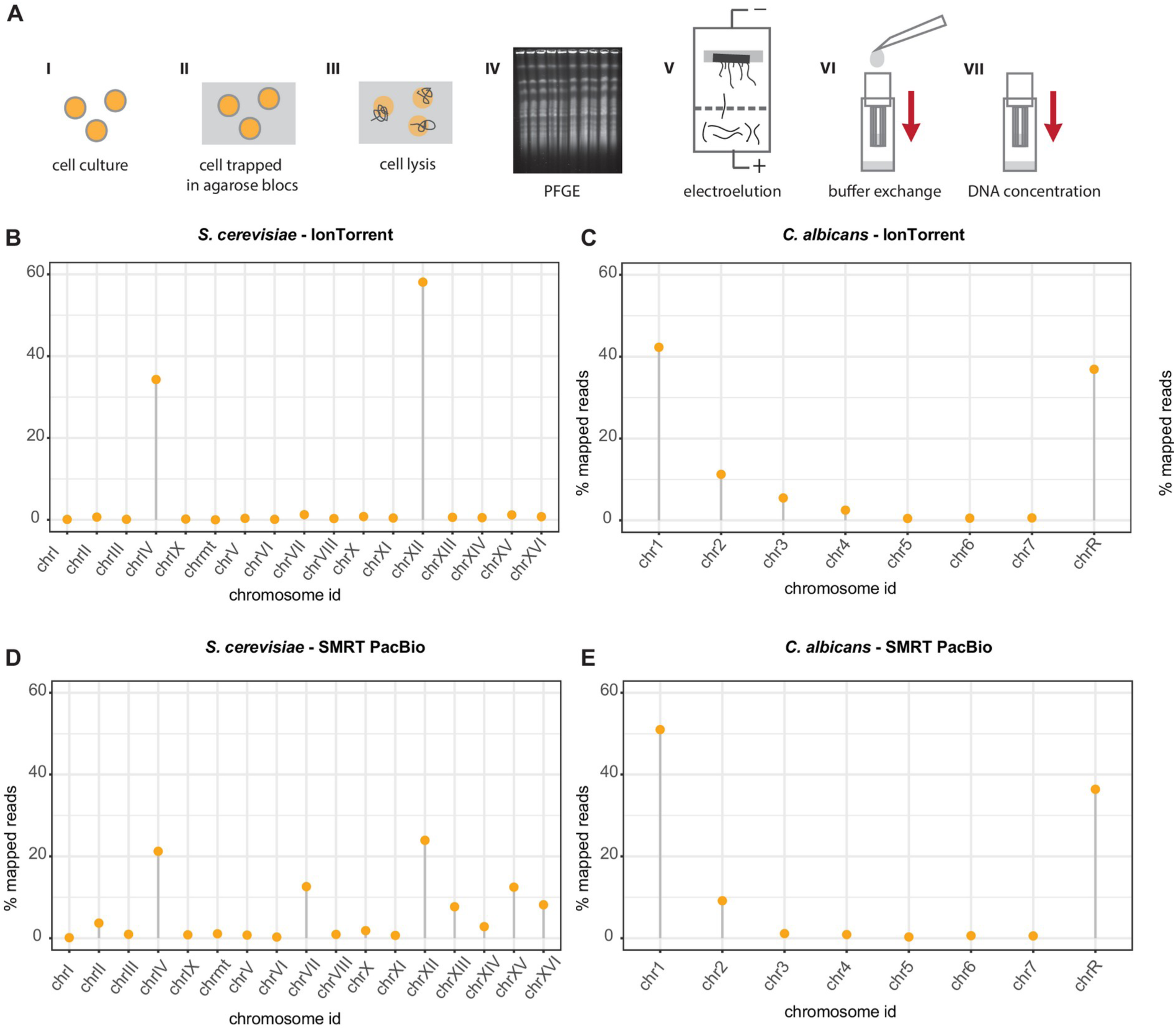
Efficiency of sequencing dataset enrichment with reads mapping to isolated chromosomes. **A** Schematic of the selective chromosome DNA isolation method. **B** *S.cerevisiae* IonTorrent dataset. **C** *C.albicans* IonTorrent dataset. **D** S. cerevisiae SMRT PacBio dataset. E C. albicans SMRT PacBio dataset.

The sequencing results obtained using both long- and short-read methods were mapped to the respective genomes. As shown in Figure 1B–E, the frequencies of reads mapped to specific chromosomes confirmed that the DNA samples were enriched for chromosomes IV and XII in *S. cerevisiae* and I and R in *C. albicans*. For long-read sequencing methods such as SMRT (PacBio) or direct DNA sequencing (ONT), the read lengths typically ranged from 20 to 500,000 bp. The use of a library size selection method enabled enrichment of the library with long DNA molecules, which in turn influenced the length distribution of the reads obtained. Furthermore, the inclusion of only selected chromosomes in the sequenced sample allowed for greater coverage of the target region using long reads.

Many eukaryotic organisms, including humans, have rDNA sequences grouped in arrays on multiple chromosomes (Sims *et al*., 2021; Rodriguez-Algarra *et al*., 2022). When total DNA is used for sequencing, accurately assigning rDNA modules to specific arrays during data analysis is a challenging task. This issue can be effectively addressed by isolating and sequencing individual chromosomes. The method we propose may be difficult to apply to organisms with rDNA arrays located on chromosomes of similar sizes. However, excluding such cases, it remains the only approach that allows for chromosome-specific reads containing repetitive regions.

Additionally, direct DNA sequencing using the ONT platform enables the enrichment of sequencing results with reads corresponding to a specific genomic region by adaptive sampling. During sequencing, decoded sequence of the initial fragment of the DNA molecule passing through the pore is mapped to the genome. If it corresponds to the target region, sequencing continues; if not, the DNA molecule is removed from the pore by reversing the polarity (Loose, Malla i Stout, 2016). However, this approach is less effective for long repeat arrays. If a sample contains total DNA from an organism with, for example, more than one rDNA array, the mapping software used by ONT can reliably assign sequences only for regions near the edges of the arrays. Reads that contain only rDNA repeats are assigned randomly during the mapping stage. Thus far, this sequencing method has mainly enabled the study of rDNA repeats located near the edges of the array. Although there have been reports of sequencing the entire rDNA array of *S. cerevisiae* in a single read, these data remain unpublished.

Finally, the ultra-long-read sequencing method requires highly pure samples with minimal fragmented DNA, which can be challenging to obtain from organisms with cell walls or environmental isolates. The method we propose does not yield DNA molecules as long as those obtained from total DNA isolation methods, such as the phenol-based approach, because the DNA molecules are exposed to tensions during electroelution. However, it offers another advantage. PFGE gel-based DNA isolation allows for the purification of DNA samples from low-molecular-weight contaminants, which is a common problem with environmental samples (Dicuonzo *et al*., 2001). Therefore, our method of DNA sample preparation for sequencing could prove useful for highly contaminated samples.

### rDNA arrays and pseudogenes

To date, no algorithm has been developed to reconstruct genomic sequences at loci characterised by long, highly similar repeats. Combining data from long-read and short-read sequencing methods also fails to resolve this issue. Using the minimap2 and miniasm algorithms, we obtained contigs containing the edges of the rDNA array and a single repeat, as well as contigs containing 2–5 consensus repeats of the modules encoding ribosomal RNA. Since the full repetitive loci sequence could not be assembled, we mapped independent sequencing datasets to the reference sequence of the chromosome containing the rDNA array: chromosome XII for S. cerevisiae and the R chromosome for C. albicans. Our aim was to estimate the number of rDNA copies using an established method based on comparing the sequencing depth of repetitive and non-repetitive genomic regions (Ganley i Kobayashi, 2007; James et al., 2016; Sharma et al., 2022). We calculated the per-base sequencing depth of the reference chromosome sequence, which enabled us to identify the repetitive regions easily (Figure 2A-D).

**Figure 2.**
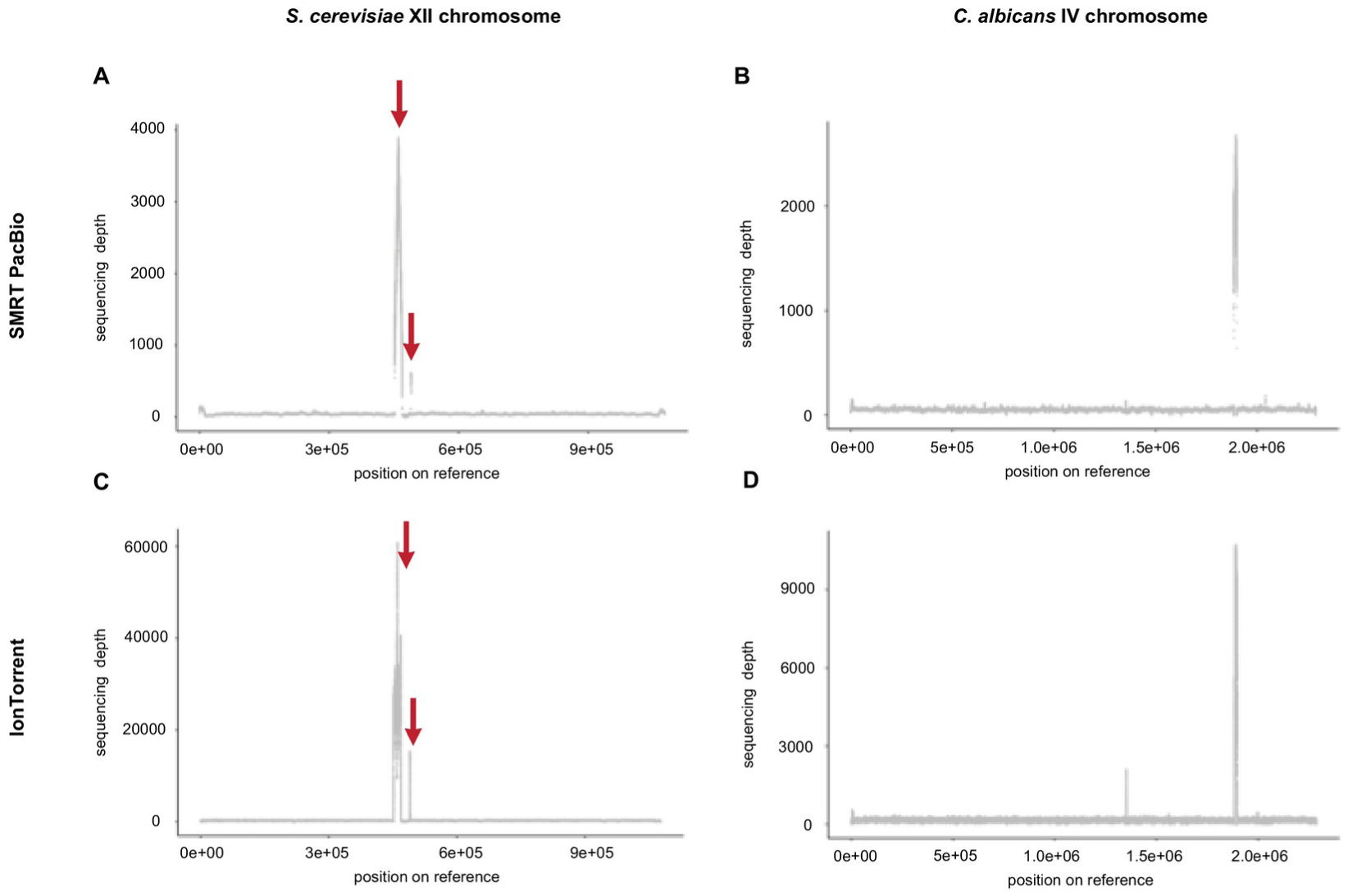
Distribution of sequencing depth across chromosomes containing rDNA arrays. **A** *S.cerevisiae* SMRT PacBio dataset mapped to reference genome, chromosome XII. **B** *C.albicans* SMRT PacBio dataset mapped to reference genome, chromosome R. **C** *S.cerevisiae* IonTorrent dataset mapped to reference genome, chromosome XII. **D** C. albicans IonTorrent dataset mapped to reference genome, chromosome R, small second peak represent RPS1.

Interestingly, in *S. cerevisiae*, two adjacent regions on chromosome XII were identified where reads containing rDNA module sequences mapped (Figure 2A and C, red arrows). The first region contains two rDNA repeats and corresponds to the location of the rDNA array on the chromosome. The second region contains a single rDNA copy encoded on the antisense strand, potentially representing a remnant of a duplication of an entire module. Short reads obtained using the IonTorrent method were randomly assigned between these two regions, highlighting the challenges posed by short-read sequencing methods. Based on sequencing depth, we estimated the number of rDNA copies in *S. cerevisiae* to be 183, consistent with similar estimates (Sharma *et al*., 2022). Reads whose lengths represent only a small fraction of the entire repetitive module may originate from pseudogene sequences but could be erroneously assigned to modules within the repeat array. In the case of *S. cerevisiae*, the presence of a single pseudogene does not significantly affect the copy number estimation. However, in humans, highly degraded but nearly full-length ribosomal DNA modules can be found at multiple sites in the genome, on chromosomes that do not contain rDNA arrays (Robicheau *et al*., 2017).

Estimating the copy number of long repeats, such as those found in rDNA *loci*, remains a challenge. Commonly used approaches including pulsed-field gel electrophoresis, droplet digital PCR (ddPCR), and whole genome sequencing unfortunately carry a degree of technical error (Morton *et al*., 2020). We demonstrated that the common method of estimating polymorphism based on the consensus alignment of all reads containing rDNA to a reference repeat sequence is flawed, as it incorporates polymorphisms from pseudogenes. This issue can be partially addressed by using long-read sequencing techniques, provided that only reads exceeding the full length of the repetitive module sequence are included. Unfortunately, both SMRT (PacBio) and ONT exhibit a higher error rate in nucleotide identification during read sequencing compared to short-read methods such as Illumina (Wang *et al*., 2012; Zhang, Jain and Aluru, 2020). Furthermore, both *C. albicans* and *S. cerevisiae* harbour rDNA copies on extrachromosomal circles (Sinclair i Guarente, 1997; Huber i Rustchenko, 2001), which complicates not only copy number estimation but also the study of rDNA polymorphism. For single nucleotide polymorphism studies, short-read sequencing methods are most commonly used (Kwan *et al*., 2013; Wang *et al*., 2020; Bizarria *et al*., 2023). However, we caution that averaging variability information at a given position within long repetitive modules can be affected by reads originating from pseudogenes, potentially leading to incorrect conclusions.

### Determination of the sequence of reference modules

Due to the lack of tools capable of assembling genomic sequences with long repeats, rDNA is typically represented by single copies, even in model organisms. To address this, we developed the rDNAmine toolkit, which bypasses the reconstruction of the rDNA matrix. Our pipeline uses long reads from ONT sequencing (Figure 3A). It can be applied to any dataset containing this type of data, provided the average read length exceeds the length of the repetitive sequence under study. For visualisation, we used two datasets from the SRA repository: total DNA sequencing of *S. cerevisiae* (Zhang *et al*., 2022) and *C. albicans* (*Microbiology Resource Announcements*, 2021). As an example of long repeats, we used rDNA modules.

**Figure 3.**
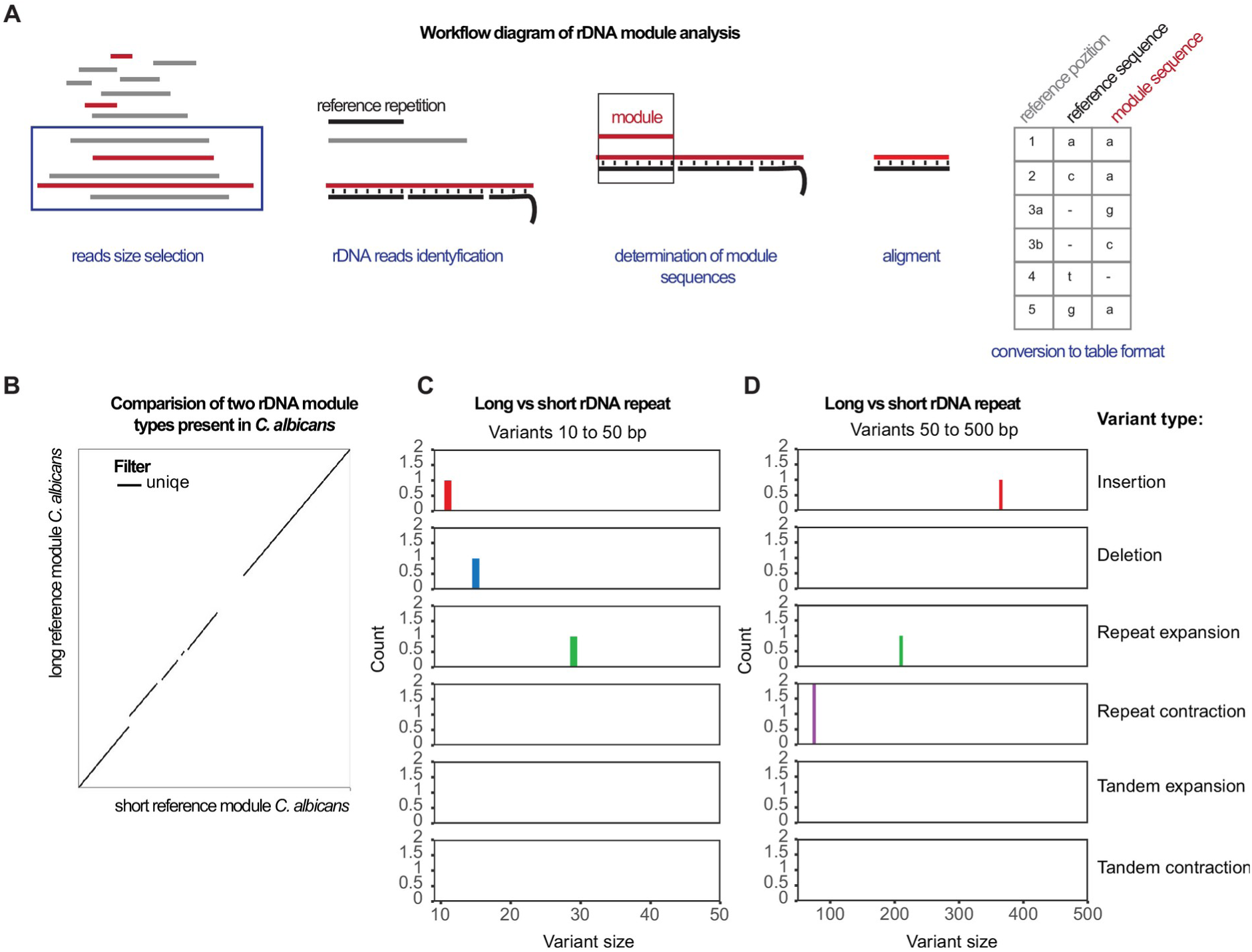
Comparison of long and short rDNA module reference sequences in C. albicans. **A** Schematic of key steps of long repetitive module isolation from ONT dataset using rDNAmine toolkit. **B** Dot plot from Assemblytics showing six classes of structural variants between long and short reference modules from the *C. albicans* ONT dataset. C Size distributions of small polymorphism events from alignments of two compared reference modules. **D** Size distributions of large polymorphism events from alignments of two compared reference modules.

The first step is to filter out reads of insufficient length. The minimum read length for analysis should exceed twice the length of the repetitive module under study. For high-quality datasets with a substantial proportion of long reads, the cut-off can be set higher. Next, the resulting pool of reads is searched using Hidden Markov Models, which map fragments of the read sequences to a specified reference sequence.

We adopted the RDN-2 sequence from the *S. cerevisiae* genome, as deposited in the SGD, as the reference sequence for the rDNA module. The HMMER algorithm outputs information about the length of the reference sequence assigned to each read, along with the precise coordinates of these assignments. A custom program then processes the output files to select, for each read containing rDNA, the modules that fall within a specified length range. Based on the selected coordinates, the module sequences are extracted from the reads and saved to separate files, preserving information about their origins. In the next step, each module is compared to the reference repeat, and these pairs are converted into tabular files with numbered reference positions. The resulting data can be easily loaded into the R environment using our custom script, rDNAmineR and analysed with any data wrangling library, such as dplyr (Wickham *et al*., 2022).

Although the sequences of rRNA-encoding genes have been determined for most described species, the full repetitive sequences present in genomes are rarely available. These can be obtained using tools such as BLAST (Camacho *et al*., 2009), which provides sequences annotated in repositories as rDNA repeats or fragments. Additionally, sequences of modules originating from complex genomes or transcriptomes may be found, but we must rely on their correct assembly. Using *C. albicans* as an example, we demonstrate how to obtain a complete rDNA module from ONT data.

For *C. albicans*, the rDNA sequence deposited in the Candida Genome Database (CGD) was too short to cover the entire repeat (Ca22chrRA_C_albicans_SC5314, 1888934-1896830). Consequently, the HMMER algorithm identified only the reads containing fragments of the reference encoding rDNA genes. Using the coordinates obtained for individual genes, we excised the full rDNA repeat sequence from the reads and used it as a reference module (Supplementary file 1). Interestingly, for some reads containing rDNA repeats, the algorithm was unable to isolate the unfragmented module. Based on the coordinates determined for the rDNA-encoding genes, we identified a second module sequence that was 464 nucleotides longer. We repeated the analyses independently using both the longer and shorter reference sequences of *C. albicans*. A comparison of the two module sequences revealed one small insertion and one large insertion in the 25S gene (Figure 3C and D). Moreover, we detected two repeat expansion events (expansion of short, few-nucleotide repeat**s)** along with one long repeat contraction (decrease in the number of short, repeated nucleotide units in a DNA sequence) (Figure 3C and D).

The large insertion present in the long modules is a group I intron located within the 25S gene, which has been used in population studies of this species (Karahan *et al*., 2004). In *C. albicans*, the length of the rDNA repetitive unit has been estimated at 12,756 nucleotides (Jones *et al*., 2004), exceeding the lengths of the modules we isolated. This earlier estimation was based on the length of a consensus sequence assembled from reads obtained via a short-read sequencing method. Our findings reflect the variation in the lengths of modules present in the long reads. The greatest differences in the lengths of *C. albicans* reference modules were observed in the ITS1 and ITS2 regions. These highly evolutionarily variable non-coding rDNA regions have been used in phylogenetic studies for many years (Montrocher *et al*., 1998; Madden *et al*., 2022). The high sequence plasticity in these intra-array regions should be considered when assessing the relatedness of closely related species.

Using an rDNA copy isolated from ONT reads as a reference, rather than an external reference, offers significant advantages when studying the internal polymorphism of rDNA arrays. rDNA sequences can vary within the array, as demonstrated in our study of *C. albicans*. Moreover, unlike pseudogenes, rDNA sequences are subject to concerted evolution and should therefore be relatively homogeneous (Ganley i Kobayashi, 2007). Determining the characteristic polymorphism landscape of the array enables comparative studies between closely related species and facilitates the identification of pseudogenes (Li *et al*., 2017). The rDNAmine tools we have described allow for the analysis of any type of repetitive sequences with long repeats.

### Characterisation of rDNA copy length polymorphism

To address the challenges of studying polymorphism in long repetitive sequences, we bypassed the mapping and assembly steps. Using rDNAmine tools, we isolated sequences of 471 long and 163 short rDNA modules from *C. albicans* and 896 rDNA modules from *S. cerevisiae*. Initially, we analysed the length distribution of rDNA modules and observed both inter- and intraspecies differences in rDNA module length distributions (Figure 4A and B).

**Figure 4.**
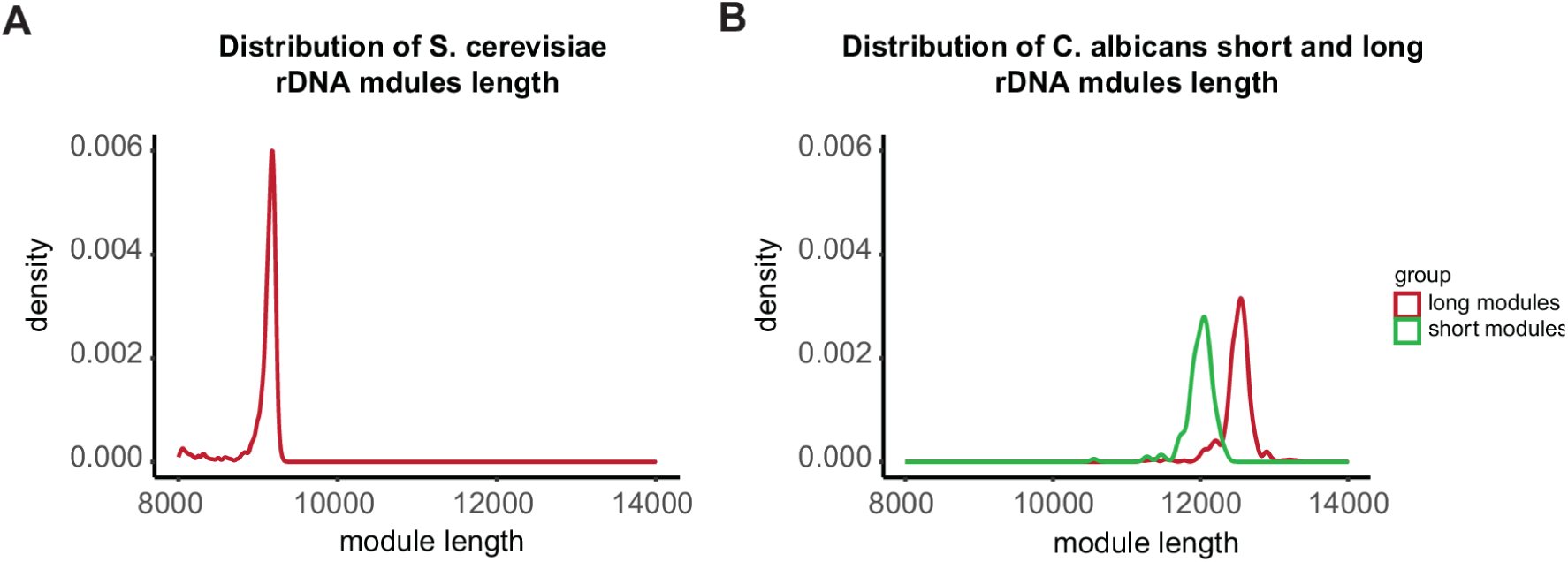
Comparison of rDNA module lengths in *S. cerevisiae* and. C. albicans**. A** Distribution of module lengths in S. cerevisiae. **B** Distribution of module lengths in the two groups of modules from C. albicans.

Modules from *S. cerevisiae* exhibited less variation in length than those from *C. albicans* (Figure 4A and B). Furthermore, short *C. albicans* modules displayed slightly greater sequence length plasticity than longer ones (Figure 4B). We believe that longer and shorter modules did not alternate in the array but instead clustered into two distinct regions. Only 8 reads contained both types of modules, superseded at their ends. Thus, it is possible that within the rDNA array on the R chromosome of *C. albicans* two distinct populations of rDNA modules form separate sub-arrays of repeats (Supplementary table 1, Supplementary file 1). Such a structure of the rDNA array corresponds with previous studies on the propagation of rDNA copies within *loci* in representatives of Ascomycetales (Ganley i Kobayashi, 2007).

### Polymorphism in *S. cerevisiae* and *C. albicans* rDNA modules

The isolated modules were required to meet minimum criteria for size and similarity to the reference module sequence, as determined by probability values estimated by the HMM algorithm (criteria are provided in the Nanopore direct DNA sequencing datasets section of Materials and Methods). This approach significantly reduces the risk of including pseudogene-derived modules in the analysis. Sequence analysis of reads obtained using ONT, conducted with our rDNAmine toolkit, enabled the creation of aligned reference-module pairs stored in a tabular format. These data were imported into the R environment and combined into a consolidated table based on the numbering of reference positions. To calculate the substitution-deletion polymorphism coefficient (SDC) only information regarding deletions and substitutions was used. SDC describes the variability of a given position or region within the rDNA repeats in the set of modules isolated from long reads. This measure can be used to compare the internal variability of the rDNA array between different species or mutants. However, it is important to note that it is not equivalent to single nucleotide polymorphism estimated based on short reads mapped to a reference. Our analysis revealed that short *C. albicans* modules and *S. cerevisiae* modules show similar distributions of SDC polymorphism rates. Most positions in the reference sequence exhibit low variability among the rDNA modules. An intriguing observation arose with the long modules of *C. albicans*. In this case, two distinct populations were evident: one with variability comparable to the short modules and another associated with modules exhibiting highly variable sequences (Figure 5A). Next, we plotted the distribution of the SDC coefficient across the entire reference module to visualise the most variable regions (Figure 5B). We observed that the greatest variability occurs in regions containing homopolymers, which is a well-known phenomenon. This results partly from the natural polymorphism of these regions and partly from the limitations of sequencing technologies and alignment algorithms.

**Figure 5.**
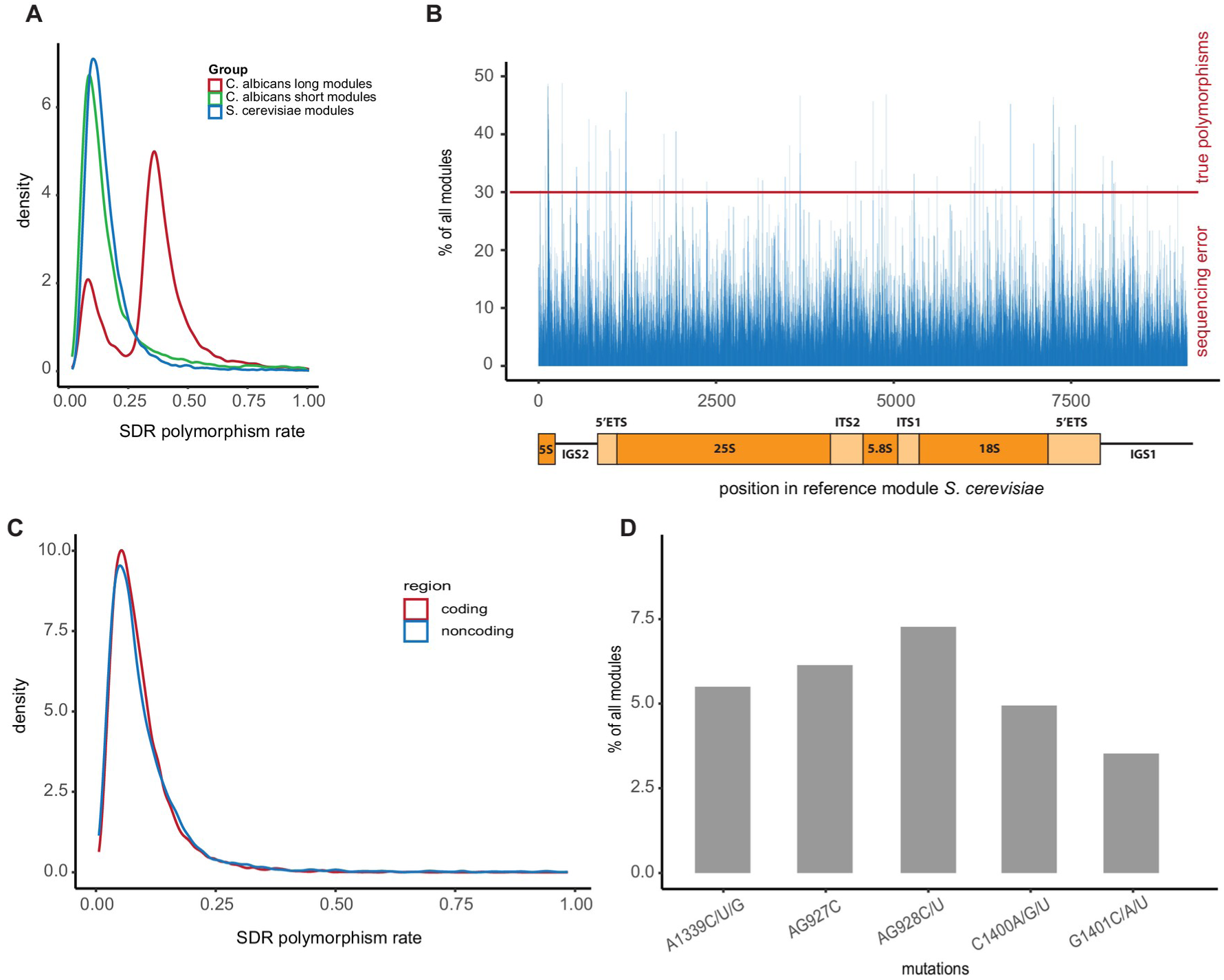
Analysis of polymorphism in rDNA modules. **A** Distribution of SDC coefficients in S. cerevisiae and C. albicans modules. **B** Distribution of SDC coefficients in coding and non-coding sequences in *S.cerevisiae* modules. Diagram of the rDNA repeat reference sequence structure in *S. cerevisiae*. **C** Frequency of rDNA modules containing polymorphism in particular base of reference. Polymorphism rates were assessed using a two-tailed Wilcoxon sum rank test with Benjamini-Hochberg correction (padj.=5.37e-08). **D** Frequency of rDNA modules containing lethal mutations.

Since the SDCs were calculated for each reference position, it was possible to compare the polymorphism rates between coding and non-coding regions of the rDNA repeat, as demonstrated for *S. cerevisiae* (Figure 5C). Polymorphism in non-coding regions was found to be statistically significantly higher than in coding regions (two-tailed Wilcoxon rank sum test with Benjamini-Hochberg correction, padj = 5.21×10⁻⁸). This result aligns with previous observations, as non-coding regions are not subjected to the same strong evolutionary conservation pressure as genes (James *et al*., 2009). Nevertheless, these results should be interpreted with caution.

In particular, sequencing accuracy must be taken into account when evaluating the observed polymorphism rates. Oxford Nanopore sequencing has an error rate of ∼5–15%, depending on the chemistry and basecalling algorithm, with systemic miscalls occurring most frequently in homopolymer regions. To minimise the impact of these technical biases, we applied a stringent threshold of 0.3, corresponding to variants detected in at least 30% of rDNA modules, above which positions were considered truly polymorphic (Figure 5A–C). Variants below this level are largely dominated by sequencing noise and alignment inaccuracies rather than genuine biological variation. Using this criterion, we estimate that roughly 85 positions within the *S. cerevisiae* rDNA module represent true polymorphic sites. This conservative approach accounts for both platform-specific limitations and alignment biases, allowing the inferred polymorphism landscape to more faithfully reflect variation within the rDNA array.

rDNA modules undergo concerted evolution, which promotes greater homogeneity among copies in the array and reduces the number of copies containing mutations that cause deleterious phenotypes (Wang *et al*., 2023). We selected six mutations in *S. cerevisiae* that are well documented in the literature as being lethal (Dong *et al*., 2008). As expected, these mutations were extremely rare, occurring in only a few hundred of the 4,854 analysed modules. This means that if the rDNA matrix contains between 150 and 200 copies of rDNA, the mutations are present in only a few (Figure 5D). The copies containing mutations at positions critical for ribosome function are likely to be transcriptionally inactive. Additionally, in the analysed population of modules extracted from long reads, we observed insertions with potential implications for the tertiary structure of rRNA molecules. However, their biological relevance will require confirmation by other methods and the exclusion of sequencing artefacts.

To improve repeat boundary determination, we specifically selected the 5S rRNA gene as the starting point of the reference repeat sequence, assuming its well-conserved evolutionary sequences would enable more precise estimation. At the other end of the repeat, a more polymorphic region, IGS1 (intergenic spacer 1), contains sequences regulating both rRNA gene expression and autonomously replicating sequences (ARS).

Given the sequencing error rates discussed above, we acknowledge that some of the rare variants detected in individual modules may represent sequencing artifacts rather than true mutations. The mutations shown in Figure 5D occurred at frequencies below our 0.3 threshold and should therefore be interpreted with caution. While the pattern of rare deleterious mutations is consistent with expectations from concerted evolution, validation using orthogonal methods would be required to confirm individual variants.

## Conclusions

The budding yeast *S. cerevisiae* is extensively used to investigate the genetic and molecular basis of eukaryotic ribosome biogenesis (Baßler i Hurt, 2019), as well as ribosome composition and function (Armache *et al*., 2010; Barandun, Hunziker i Klinge, 2018; Fujii *et al*., 2018). It also serves as a valuable model for examining rDNA polymorphism. Due to the structure of rDNA *loci*, previous studies on the organisation rRNA-encoding genes and the architecture of ribosomal active sites relied on an *S. cerevisiae* strain in which the entire rDNA array was deleted and replaced with a single functional rDNA copy introduced on a high-copy plasmid (Wai *et al*., 2000).

Here, we present novel tools for analysing repetitive sequences characterised by long tandem modules. Our selective chromosome isolation method enables extraction of high-quality DNA suitable for both short- and long-read sequencing. The resulting dataset comprises exclusively repeats from a specific chromosome, allowing precise investigation of copy variability within a given genomic *locus*. For rDNA studies in particular, the polymorphism profiles obtained are unaffected by circular DNA molecules containing rDNA copies (rDNA loops), which are commonly found in eukaryotic genomes (Møller *et al*., 2015; D’Alfonso, Micheli i Camilloni, 2024; Gumińska *et al*., 2024).

We additionally introduce an efficient strategy for analysing long reads containing extended repeats. This approach eliminates the need for global alignment of all repeated modules to a reference, substantially reducing processing time and simplifying downstream workflows. Our toolkit, rDNAmine, identifies and isolates sequences harbouring long repeats from reads generated via ONT direct DNA sequencing. Individual modules are aligned to a reference and stored in a tabular format, facilitating straightforward analysis in R. Notably, this framework allows detection of polymorphisms co-occurring within a single repeat unit.

The reliability of the obtained module sequences depends on the intrinsic error rate of the long-read sequencing technology employed. In the case of Oxford Nanopore sequencing, the standard sequencing protocol yields accuracies of 97.6% (R9.4.1) and 99.1% (R10.4 simplex). These values are influenced by the chemistry used, the generation of the flow cell, and the basecalling algorithm, which continues to improve with successive software update. Data generated using this approach are suitable for the investigation of long repeat structural polymorphisms, as we showed for *C. albicans* rDNA. However, they remain insufficient for accurate SNP calling. For studies focused on single-nucleotide mutations or SNP detection, we recommend the Oxford Nanopore duplex system, which currently achieves an accuracy of 99.7–99.9% (R10.4 duplex) (Kim et al., 2025; Stoeck et al., 2024).

Whereas most alignment-based polymorphism analysis tools are limited by sequence number or length, rDNAmine overcomes these limitations. Although we used MAFFT for alignment, any other pairwise aligner accepting FASTA input can be substituted.

rDNA modules served as a well-characterised model system to demonstrate performance on real Oxford Nanopore datasets, which are inherently noisy. We did not benchmark the method on simulated data, focusing instead on applicability to experimentally generated reads. The large-molecule enrichment and bioinformatic screening steps outlined here represent initial stages of a broader analytical workflow and are not intended to directly address biological questions. They rather provide a means to obtain enriched material and identify candidate reads for downstream investigation. Consequently, biological interpretation must be undertaken in the context of specific experimental designs, probability thresholds should be applied stringently, and all findings validated using orthogonal methods. We anticipate that our proposed methods will advance research on repetitive genomic regions. rDNAmine is particularly well suited to investigating length variability within long repeat arrays, as demonstrated for *C. albicans*. For SNP-focused studies, a long-read sequencing method with a very low error rate, such as the Oxford Nanopore duplex system, is required.

## Acknowledgements

We thank Andrzej Dziembowski for the assistance in preparing this manuscript and for providing computational resources essential for bioinformatics analyses. We also thank Mariusz Czarnocki-Cieciura for his support. This research was funded by the Polish National Science Center 2016/23/N/NZ2/01163 to AC.

## Contribution

ACC was responsible for writing and testing the rDNAmine bioinformatics tools, isolating DNA material, sequencing, performing bioinformatics analyses of all datasets, and preparing the figures. NG was responsible for testing the bioinformatics tools, creating the GitHub repository, preparing the graphical abstract, and figures. ACC and NG wrote and revised the manuscript.

