## Supplementary figures and images for "rDNAmine: A New Tool for the Analysis of Long Repetitive Sequences"

### SF1_BioRxiv.pdf

**Supplementary Figure 1.**

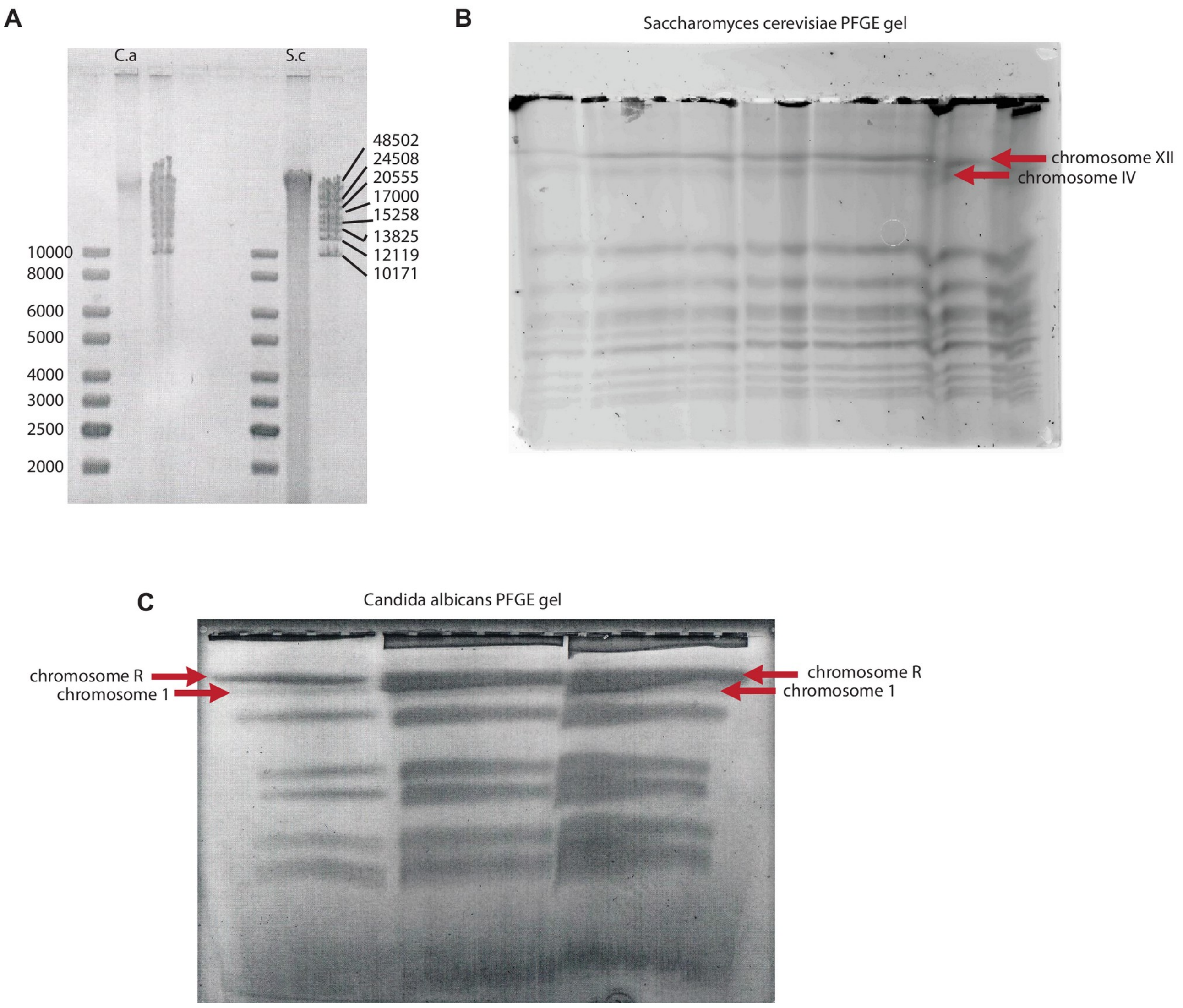
